# Dynamics of species-rich predator–prey networks and seasonal alternations of keystone species

**DOI:** 10.1101/2022.10.16.512452

**Authors:** Sayaka Suzuki, Yuki G. Baba, Hirokazu Toju

**Affiliations:** Center for Ecological Research, Kyoto University, Otsu, Shiga 520-2133, Japan; Institute for Agro-Environmental Sciences, NARO, Kannondai 3-1-3, Tsukuba, Ibaraki 305-8604, Japan

## Abstract

In nature, entangled webs of predator–prey interactions constitute the backbones of ecosystems. Uncovering the network architecture of such trophic interactions has been recognized as the essential step for exploring species with great impacts on ecosystem-level phenomena and functions. However, it has remained a major challenge to reveal how species-rich networks of predator–prey interactions are continually reshaped though time in the wild. We here show that dynamics of species-rich predator–prey interactions can be characterized by remarkable network structural changes and alternations of potential keystone species. Based on high-throughput detection of prey DNA from 1,556 spider individuals collected in a grassland ecosystem, we reconstructed dynamics of interaction networks involving, in total, 50 spider species and 974 prey species/strains through eight months. The networks were compartmentalized into modules (groups) of closely interacting predators and prey in some but not all months. As the modules differed in detritus/grazing food chain properties, complex fission-fusion dynamics of below-ground and above-ground energy channels was reconstructed across the seasons. The substantial shifts of network structure entailed alternations of spider species located at the core positions within the entangled webs of interactions. These results indicate that knowledge of dynamically shifting food webs is essential for understanding the temporally-varying roles of keystone species that interlink multiple energy channels.

---

Predator–prey interactions are among the most ubiquitous types of interspecific interactions in terrestrial and aquatic ecosystems^1–5^. A number of classic studies have discussed how structure of predator–prey interaction networks (i.e., food webs) underlies ecosystem-level characteristics and phenomena^6–9^. For instance, the presence of generalist predators in food webs may lead to community- or ecosystem-level stability because such predators suppress populations of many lower-trophic consumers, thereby preventing outbreak events^9,10^ (i.e., top-down effects). In general, information of predator–prey network structure provides bird’s-eye views of energy or material flows in ecosystem-level processes^11–14^, allowing us to highlight specific species or interactions that have profound effects on stability and functions of ecosystems^15,16^.

Nonetheless, we still have limited knowledge about how predator–prey network structure *per se* changes through time^2,17–19^. Phenological dynamics and migration of species, for example, cause turnover of species compositions within a community, inevitably altering structure of interaction networks^20–22^. Moreover, even if predator and prey community compositions are unchanged, the nature of interactions can change through time, entailing network rewiring^22,23^. Despite the potentially dynamic nature of predator–prey systems, majority of the existing data of consumer–victim interactions are “snap-shot” records of interactions observed within short time windows or “cyclopedic” records of all possible interactions that can occur in long time periods^22,24^. Although there have been pioneering studies reporting temporal dynamics of interactions between guilds or trophic groups^25^, time-series data with species-level resolution remain scarce.

If available, detailed information about dynamics of species-rich predator–prey systems will provide platforms for understanding mechanisms by which community-level stability/instability is determined in nature^14,26,27^. Such information, for example, is expected to help us investigate how multiple energy channels, such as those deriving from detritus and grazing food chains, are dynamically integrated or disconnected in ecosystem processes^11^. Moreover, it will allow us to examine the potential that keystone species^9,10^ are altered through dynamics of predator–prey interactions.

Here we report how predator–prey interaction networks consisting of hundreds of species are dynamically reshaped through time. We conducted intensive field sampling of spider communities in a temperate grassland ecosystem in Japan and then analyzed Hexapoda prey DNA for each of the predator (spider) specimens collected across eight time points from April to November. The rich dataset composed of 50 predator (spider) species and 974 prey operational taxonomic units (OTUs) allowed us to reveal how predator–prey interaction webs could be compartmentalized into substructures (network modules^14,28^) that differed in grazing/detritus food chain properties. We then tested the hypothesis that potential keystone species within the predator–prey systems could be altered through the time-series. By compiling the network data of the eight time points, we further highlighted potential keystone species that linked above- and below-ground energy channels across the seasons.

## Results

### Predator and prey diversity

We randomly collected 168–441 spider individuals per month with sweeping nets in a temperate grassland from April to November, 2018. In total, 2,224 spider specimens representing 63 species (35 genera and 15 families) were collected. The species compositions of the spider community changed dramatically through seasons (Fig. 1a).

**Fig. 1.**
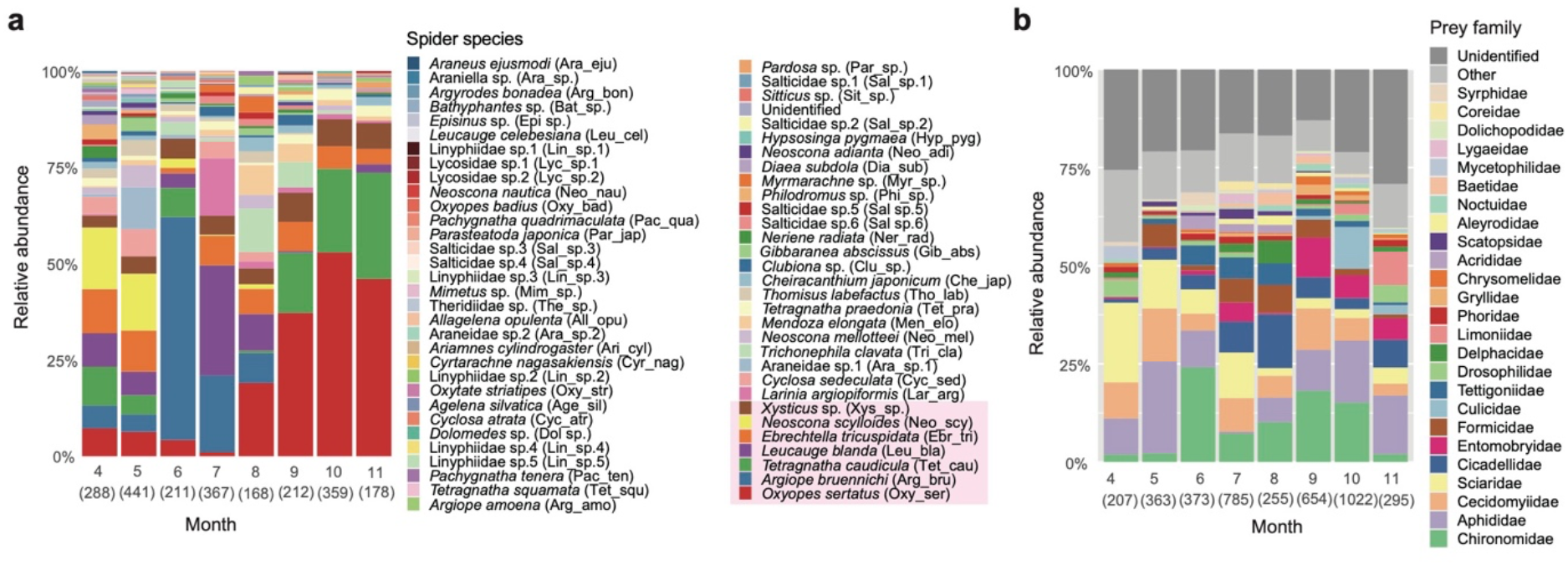
Predator and prey diversity. **a**, Spider species composition. The composition of collected spider species is shown for each month with total number of individuals in a parenthesis. Abbreviations of species names used in Figure 2 are shown in parentheses. The seven most abundant species in the dataset are highlighted. **b**, Prey taxonomy. The composition of prey detection counts is shown with family-level taxonomy. The total number of prey detection counts is shown in a parenthesis for each month. See Extended Data Figure 1 for genus- and order-level taxonomy of prey.

Based on the DNA metabarcoding of respective specimens, prey DNA was detected from 1,556 out of the 2,224 (70.0 %) collected sider individuals representing 50 species. In the dataset of each month, a prey OTU was designated as present based on a threshold number of sequencing reads (see Methods), yielding an “occurrence” matrix representing the presence (1) and absence (0) of each prey OTU in each spider sample (individual) in each month. On average, 3.34 prey OTUs were observed from single spider individuals. Based on the binary sample-level matrix of each month, we obtained a “species-level” matrix, in which rows represented spider species, columns depicted Hexapoda OTUs, and cell entries indicated the number of samples from which respective spider–Hexapoda OTU combinations were observed (hereafter, prey detection count). In total, 974 prey OTUs (defined with 97 % threshold identity) belonging to 120 families (17 orders and 3 classes) were detected from the entire dataset (Fig. 1b). The total number of predator–prey links observed across the eight months was 2,247, representing 5,190 prey detection counts.

The taxonomic compositions of the detected prey shifted considerably through the seasons (Fig. 1b; Extended Data Figs. 1-2). The prey compositions from April to May were characterized by the high proportion of Sciaridae (dark-winged fungus gnats), which have been known as a major component of detritivore Hexapoda communities^29^. In June, Chironomidae (nonbiting midges), another major detritivore taxon, suddenly became dominant within the prey contents, occupying a relatively high share until October. From July to September, a herbivore family Cicadellidae (leaf hoppers) and a predator family Formicidae (ants) appeared at high proportions in the prey data. The detritivore family Entomobryidae (slender springtails) was one of the major prey components in July, September, October, and November. Aphididae (aphids) and Cecidomyiidae (gall midges) were detected at high proportions throughout the seasons. When the Hexapoda families were roughly classified into guilds representing their major diets, both detritivores and herbivores had high shares of prey components throughout the seasons (Extended Data Fig. 3). Predator/parasite taxa were detected as well from spring (April) to late autumn (November), although their proportions in the prey detection counts was smaller than that of detritivores and herbivores (Extended Data Fig. 3).

### Predator–prey network dynamics

The network structure of spider–prey interactions changed remarkably through time (Fig. 2; Extended Data Fig. 4). The proportion of observed (realized) predator–prey interactions to all possible interactions (i.e., connectance) remained below 0.1 from April to September, while it increased in late autumn (October and November; Fig. 2b). The community-level specificity^30^ of the predator–prey interactions was significantly higher than that expected by chance in all the months, although the level of specificity dropped in June and November (Fig. 2c). The network nestedness^31,32^, which represented the extent to which specialist species interact with subsets of enemies/partners of generalist species, was significantly low (i.e., “anti-nested^32,33^”) only in some months (Fig. 2d). Meanwhile, the degree to which the networks were compartmentalized into modules of closely associated predators and prey^14,28^ (i.e., modularity^34^) showed temporal dynamics similar to those of community-level specificity of the predator–prey interactions. Specifically, network modularity was very high (standardized modularity > 3) in spring (April and May) and early summer (July), while it dropped in June and autumn (from September to November; Fig. 2e).

**Fig. 2.**
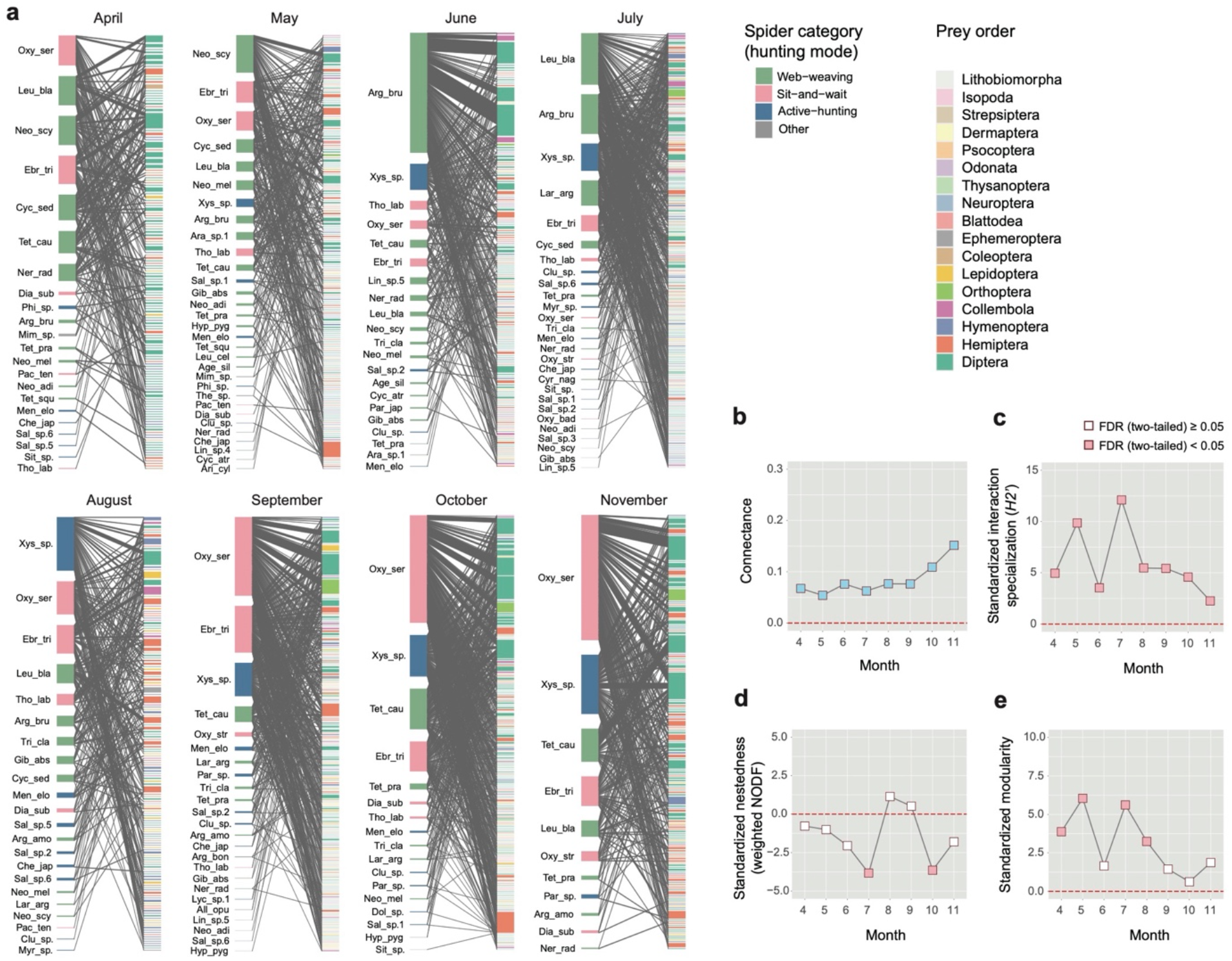
Predator–prey network dynamics. **a**, Bipartite networks predator–prey interactions. Within the network of each month, a link (edges) represents detection of a prey OTU from a spider species. The size of predator/prey vertices is proportional to prey detection counts. The hunting type of spiders and the order-level taxonomy of prey are represented by colors. **b**, Dynamics of network connectance. Changes in connectance (i.e., proportion of observed interactions to all possible interactions) are shown through the months. **c**, *H2’* metric of community-level interaction specialization. The values are standardized based on a randomization analysis. Positive/negative values indicate that network index estimates of observed predator–prey interaction matrices are larger/smaller than those of randomized matrices. **d**, Weighted NODF metric of nestedness. **e**, Barber’s modularity index of network compartmentalization.

The analysis of the succession of network modules indicated that the spider–prey interactions had been dynamically reshaped through the seasons (Fig. 3a). The network modules detected in respective months (Extended Data Fig. 5) varied considerably in their compositions of prey guilds (Fig. 3b). Such modules differing in the proportions of detritivorous/herbivorous prey were frequently merged into single modules in subsequent months through the seasons (Fig. 3a). For example, a detritivorous-prey-dominated module represented by *Argiope bruennichi* (Araneidae) and many small modules observed in May were integrated into a large module including diverse prey and spiders in June (Module June-1; Fig. 3; Extended Data Figs. 4-5). The large module observed in June was then dismantled into several modules differing in prey guild compositions in July. The merger and demolition of network modules occurred again around October (Fig. 3).

**Fig. 3.**
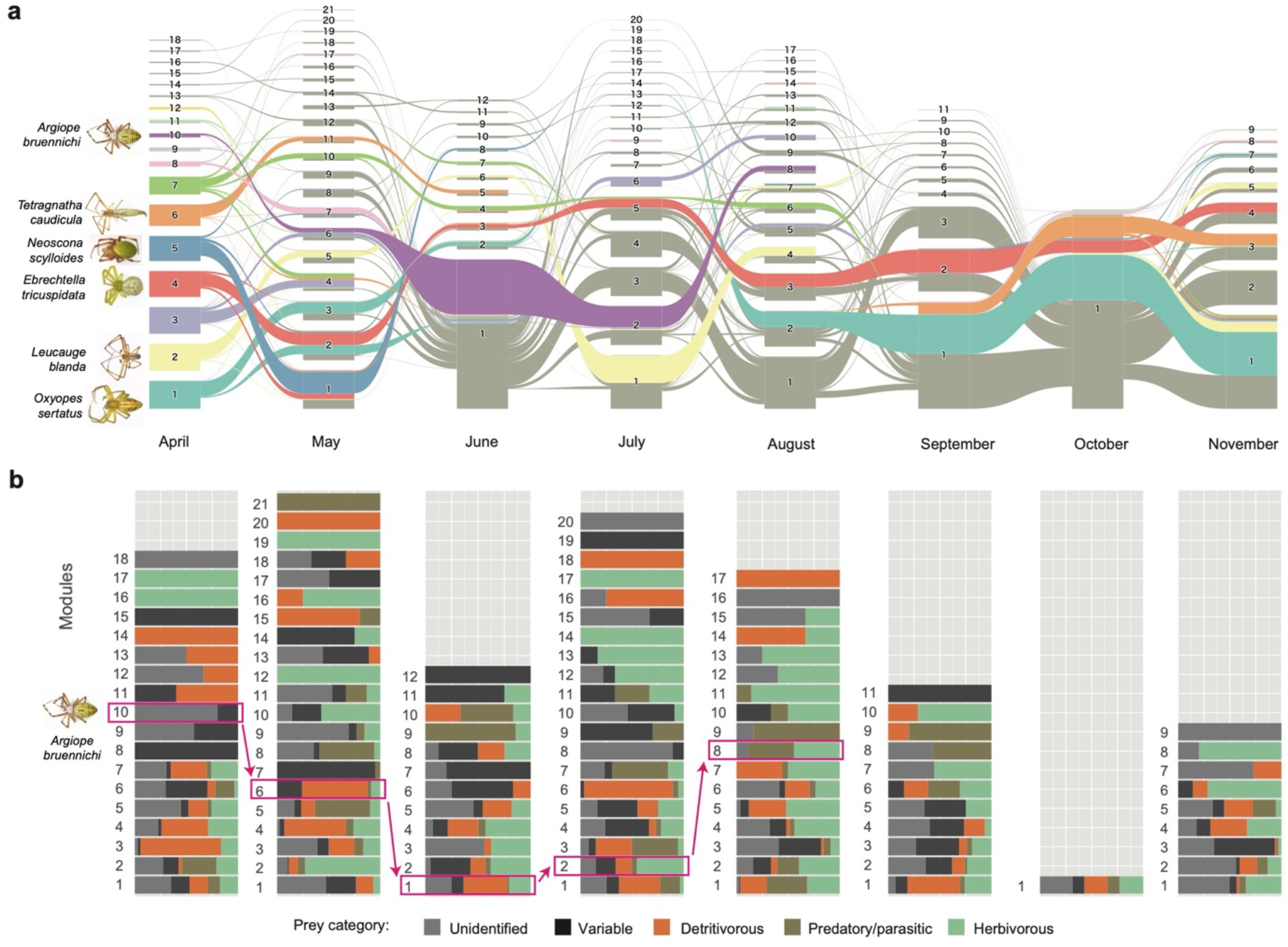
Succession of network modules. **a**, Flow diagram of network modules. The color indicates how the components of modules (spider species and prey OTUs) at each month derived from the modules defined in April. See Extended Data Figure 5 for detailed information of the inferred modules. The size of modules is proportional to the number of spider species and prey OTUs included in the modules. The network modules that included the six most frequently observed species of spiders are indicated. Among the seven most common spider species highlighted in Figure 1a, six appeared from April. **b**, Properties of network modules. For each network module observed in each month, the composition of prey detection counts is shown in terms of putative ecological guilds of prey. The flow of the modules including *A. bruennichi* across the seasons (see panel **a** for details) is highlighted.

### Alternations of keystone species

Given the high temporal variability of the spider–prey network structure (Figs. 2-3), we next examined whether potential keystone species could be altered through the seasons. In inferring the “keystoneness” of species/OTUs based on the data, we used two types of metrics. One was a standardized index of specificity to enemies/partners (standardized Kullback-Leibler distance^30^; *d’*). As the *d’* index, by definition, varies from 0 (minimum specificity given a species abundance within the community) to 1 (maximum specificity given a species abundance), the value 1 – *d’* was used to evaluate the extent to which a spider species or prey OTU interacted with broad ranges of prey/predators (hereafter, “interaction generality”). The other metric was betweenness network centrality^35^, which was a measure of the degree to which a focal network vertex is located within the shortest paths interconnecting pairs of vertices within a network. Because species located at the central positions within networks are expected to influence the population dynamics of many other species within the community^9,15^, the centrality metric was used to explore potential keystone predators and prey.

We then found that spider species with highest interaction generality and betweenness network centrality shifted across the seasons (Fig. 4). While many prey OTUs displayed high interaction generality, only some spider species had normalized betweenness scores greater than 0.25 (i.e., working as potential mediators of more than one fourth of vertex pairs; Fig. 4). Specifically, while *Neoscona scylloides* (Araneidae) had been the spider species with the highest interaction generality and betweenness centrality in April and May, *A. bruennichi* showed emergently high betweenness in June (normalized betweenness = 0.6657; Fig. 4). In July, *A. bruennichi* continued to have high betweenness, while another species, *Leucauge blanda* (Tetragnathidae), started to show high betweenness (Fig. 4). In August, *Xysticus* sp. (Thomisidae), *Oxyopes sertatus* (Oxyopidae), and *Ebrechtella tricuspidate* (Thomisidae) displayed moderately high betweenness. From September to November, *O. sertatus* showed the highest betweenness within the community (Fig. 4).

**Fig. 4.**
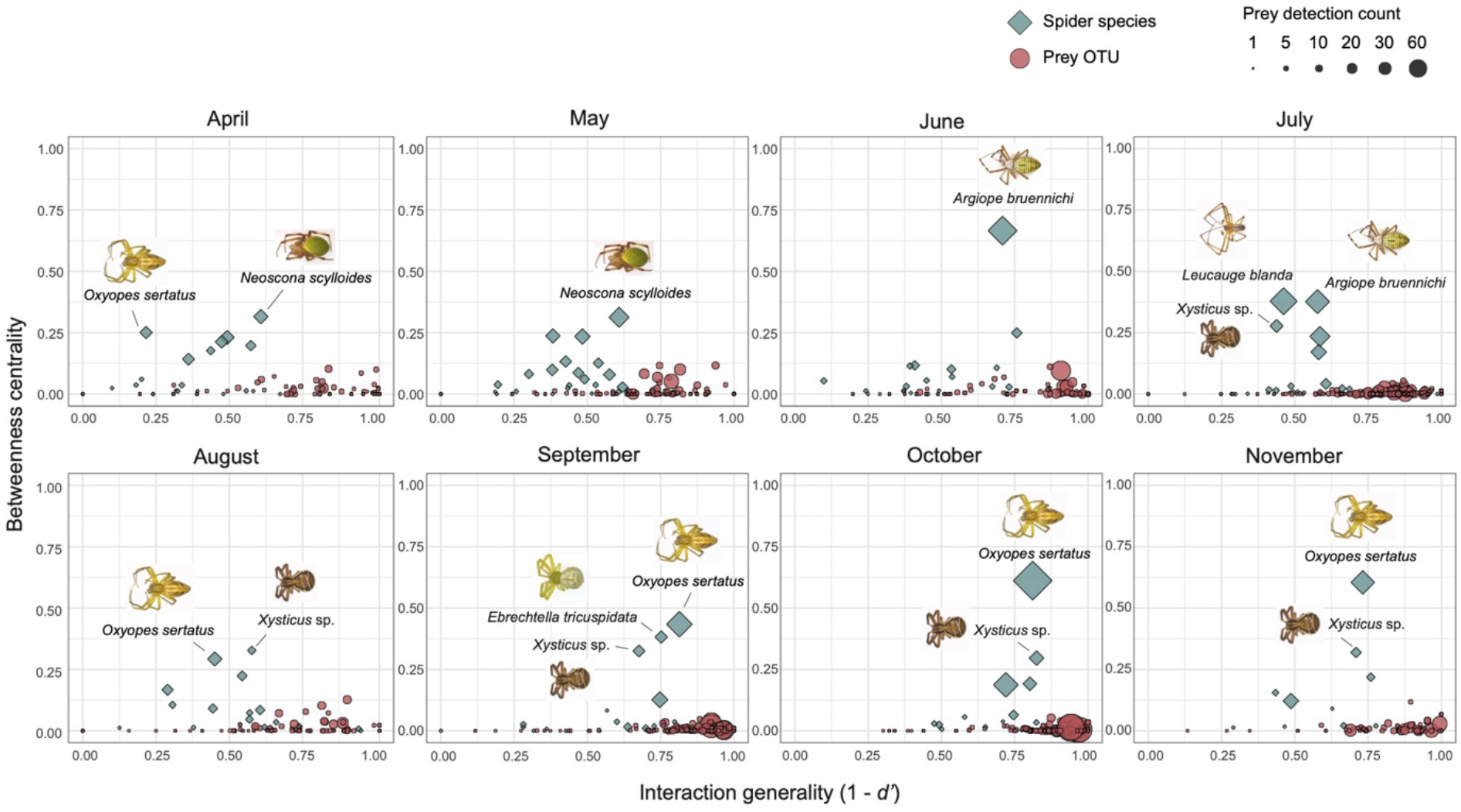
Seasonal alternations of potential keystone species. For each spider species and prey OTU in each month, the *d’* index of standardized Kullback-Leibler distance was calculated as a measure of interaction specificity. The interaction generality of each species/OTU was then evaluated as 1 – *d’* as shown along the horizontal axes. Likewise, betweenness centrality within each network was calculated for each species/OTU as shown along the vertical axes. A high value of betweenness indicates that a species/OTU is located at the central positions connecting pairs of other species/OTUs within a target network. Both indices were normalized to vary between 0 to 1. Spider species whose betweenness centrality exceed 0.25 are indicated with their names.

We further examined how interaction generality and betweenness centrality could vary within the time-series of each spider species. In our data, three spider species were present in seven or eight months from spring to late autumn. Among them, *O. sertatus* showed the highest interaction generality and betweenness centrality in the months in which it dominated the spider community (from September to November; Fig. 1a; Extended Data Fig. 6). Meanwhile, *E. tricuspidata* and *Xysticus* sp. did not necessarily display highest interaction generality or betweenness centrality in the months in which they were abundant (Extended Data Fig. 6).

### Keystones across the seasons

We next explored potential keystone species within the “meta-network” or “meta-web^22^” representing all the interactions observed from April to November (Fig. 5a). A network modularity analysis indicated that the meta-network could be compartmentalized into several modules. We then evaluated respective species/OTUs in terms of their topological roles within and across the modules. The analysis indicated that some spider species preying on diverse prey OTUs (i.e., species with high within-module degree) were placed at the topological positions interlinking multiple modules within the meta-network as represented by high among-module connectivity scores^36,37^ (Fig. 5b). When vertices within the network were classified with the two indices based on a previously proposed criteria^36^, no prey were designated as “super generalists” (Fig. 5b).

**Fig. 5.**
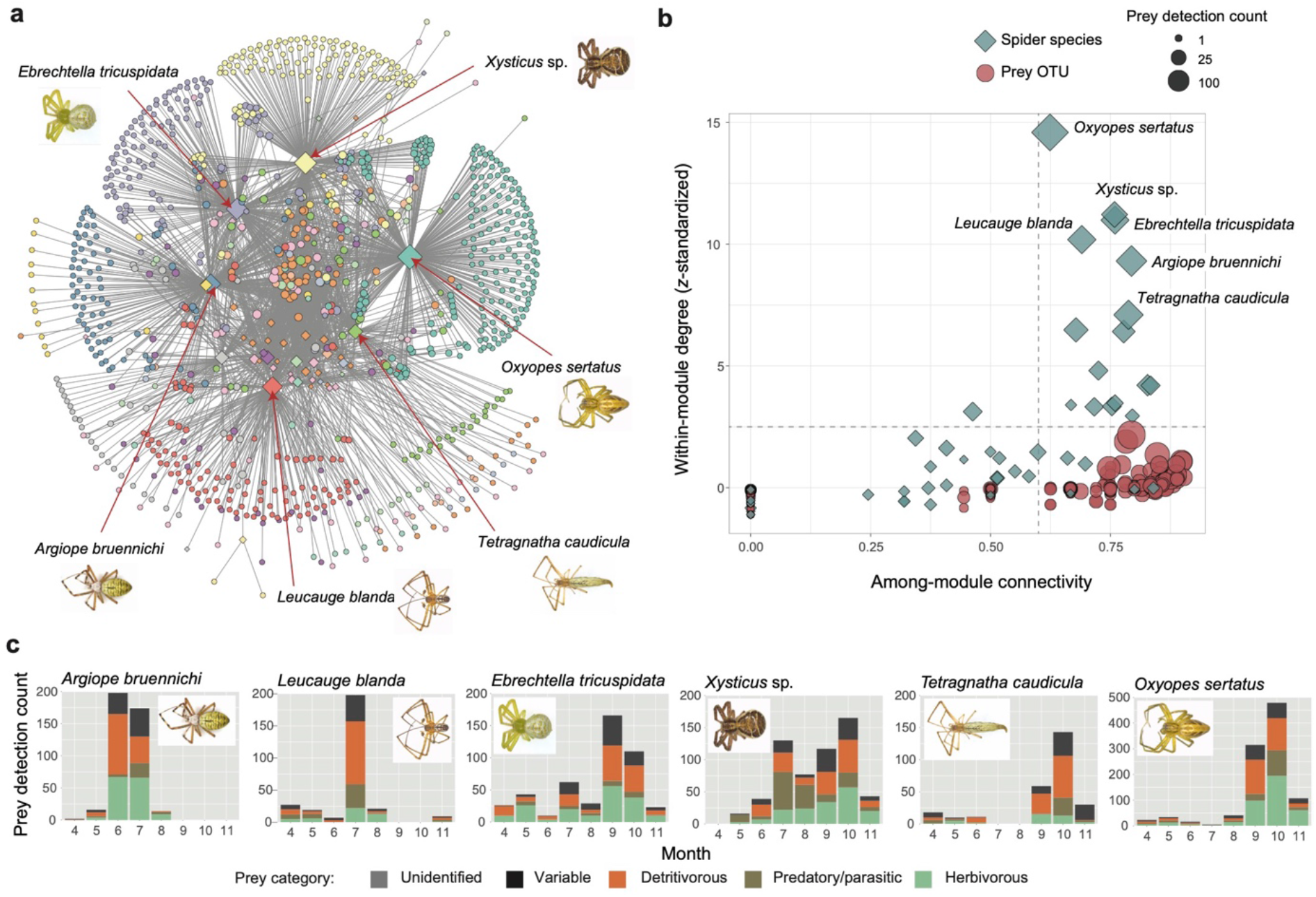
Potential keystone species across the seasons. **a**, Meta-network depicting all the predator–prey interactions observed from April to November. For each module within the meta-network, compositions (proportions) of prey detection counts are indicated in terms of month, family-level taxonomy of prey, and ecological guilds or prey. **b**, Topological roles within the meta-network. For each species/OTU within the meta-network, *z-*standardized within-module degree and among-module connectivity are shown. The lines indicate previously proposed thresholds for designating “super generalists” in ecological networks^36^. **c**, Phenology of the spider species highlighted within the meta-network. For spider species simultaneously showing high within-module degree and high among-module connectivity (panel **b**), temporal dynamics of prey detection counts are shown.

Among the spider species classified as super generalists^36^ in terms of their high within-module degree (> 2.5) and high among-module connectivity (> 0.62), *A. bruennichi* preyed on both detritivores and herbivores as a dominant predator in June and July (Fig. 5c). Albeit its sharp peak of occurrence, *L. blanda* temporally interlinked the predator–prey interactions in the transition from dominance by *A. bruennichi* to the subsequent regime of *O. sertatus* (Figs. 1a and 5c). In contrast, *E. tricuspidata* and *Xysticus* sp. were characterized as their long-time participation in the food web across the seasons (Fig. 5c). From September to November, *O. sertatus* had preyed on both detritivores and herbivores at high proportions, while the diet contents of *Tetragnatha caudicula* (Tetragnathidae) was biased towards detritivores (Fig. 5c).

## Discussion

We here showed how structure of predator–prey interaction networks could dynamically change through time. Although dynamics of predator–prey interactions themselves have been the target of intensive theoretical investigations^38,39^, empirical studies uncovering dynamics of entangled webs of interactions in species-rich predator–prey systems have remained scarce. We here explored the frontier of predator–prey interaction studies based on a high-throughput DNA metabarcoding analysis of more than 2,000 predator individuals. Our analysis indicated how tens of predator species and hundreds of prey species/strains could form webs of interactions over eight time points, providing one of the most detailed datasets of consumer– victim interactions in the history of community ecology.

The reconstructed spider–prey interactions exhibited complex community-scale dynamics characterized by the fission and fusion of network modules consisting of closely interacting species. Because those network modules differed in their prey-guild compositions (detritivores, herbivores, or predators/parasites), the results suggested the occurrence of biomass flow between different types of food chains or energy channels. While there have been a number of empirical studies supporting the concept of such subsidy effects^40–42^, our species-level-resolution data provided platforms for tracking detailed dynamics of material flow at the intersection of food chains.

The linkage of detritus and grazing food chains is of particular interest in terms of the stability of terrestrial ecosystems. While large amounts of biomass flow into detritus food chains in the forms of dead plant materials and rewards to mycorrhizal fungi^43^, only a fraction of net primary production is directly incorporated into grazing food chains^44^. Therefore, detritus-derived flow of biomass has been hypothesized to increase predator (spider) biomass disproportionately to the biomass of herbivores, imposing strong top-down control in above-ground ecosystems^44^. Our data suggested that such integration of below-ground energy channels into above-ground ones can occur continually through the dynamics of predator– prey interactions from spring to late autumn. Further empirical studies on entangled interactions at the interface of below-ground and above-ground communities^43,45^ are essential for understanding the reason why outbreaks of herbivores are frequently observed in some types of ecosystems (e.g., savannas in which outbreaks of desert locusts take place) but not in others (e.g., tropical rain forests).

The time-series data of species-rich interaction networks also provided a novel opportunity to uncover the succession of species placed at the core of predator–prey interaction networks (Fig. 5). This result indicates that potential impacts of a species to the entire communities cannot be inferred without specifying time scales and timing. In other words, even if potential keystone species are highlighted within a “meta-network” depicting all possible interactions, their impacts on community-level phenomena can be limited within short time periods (Fig. 5). Therefore, more feedback between empirical and theoretical studies is necessary for understanding dynamic nature of species’ contributions to community-level processes. Experimental exclusion of a target predator species^5^ at different seasons, for example, will enhance our knowledge about temporally-varying roles of potential keystone species within ecosystems.

For further understanding the dynamics of species-rich communities in the wild, studies on predator–prey interactions need to be extended to several directions. The processes behind the succession of potential keystone species, for example, deserve intensive research. Although species turnover through seasons imposes baseline effects, changes in species interactions can be major factors of such succession^22^. Prey switching through the ontogeny of predators^46^ is of particular interest in terms of potential mechanisms by which flexibility of interaction networks contribute to the stability of communities^23^. Extending the target of research towards more comprehensive webs of interactions is another fruitful direction of future studies. Although an estimated 400–800 million tons of prey are annually killed by spiders on the Earth^47^, spider–prey interactions are no more than subsets of terrestrial food webs. Therefore, ecosystem-level significance of the belowground–aboveground linkage connected by spiders may be further highlighted by taking into account external biological community components^48^ such as plant–herbivore and spider–vertebrate-predator interactions. Time-series data of species-rich webs of interactions will reorganize our knowledge about the stability of ecosystem-level processes.

## Methods

### Study site and sampling

Fieldwork was performed in the warm-temperate grassland located at Center for Ecological Research, Kyoto University, Shiga Prefecture, Japan (34°58’16.7”N 135°57’32.3”E). The grassland was located at the boundary of an oak-dominated secondary forest and an experimental farm. The average annual temperature and annual rainfall at the study site was 14.97 °C and 1,755 mm, respectively. The vegetation of the study site was mainly constituted by *Imperata cylindrica* (Poaceae), *Solidago canadensis* (Asteraceae), and *Artemisia princeps* (Asteraceae), while diverse taxonomic groups of herbaceous plants occurred as well. Spiders were randomly collected by sweeping with an insect net (diameter = 50 cm) on 3–5 days in the middle of each month from April to November, 2018. Owing to this sampling method, the dataset of each month analyzed in this study (see below) represented food-web structure within relatively short time periods. All spiders greater than 2 mm in body length were collected individually in 2-mL microtubes (132-620C; Watson, Fukaekasei, Kobe) or 15-mL centrifuge tubes (188271-013; Greiner Bio-One, Kremsmünster), immediately placed in a cooler container. The samples were stored at -80 °C in the laboratory. All spiders were identified to species based on morphology. Ontogenetic stages of spiders (juveniles or adults) were recorded in this identification process. In total, 2,224 spider specimens representing 63 species were collected.

### Preparation of sample and DNA extraction

To remove any DNA attached to the surfaces of spider bodies, each sample was washed through several steps. Specifically, each spider sample immersed in 70 % ethanol in a 2-mL microtube was cleaned using an ultrasonic cleaner (AS482; ASONE, Osaka) for 3 min. The samples were further washed sequentially with 1.2 ml distilled water, 70 % ethanol, and 100 % ethanol. These washing steps allowed us to analyze prey DNA that remained inside spider bodies.

After washing, each sample was ground with a homogenizer pestle (1005-39; ASONE, Osaka) in 500–700 μl of lysis buffer in a 1.5 mL tube (0.05% SDS, 20 mM Tris, 2.5 mM EDTA, 0.4 M NaCl). The sample solutions were mixed using a shaker (TWIN 3-28N; SCINICS, Tokyo) for 30 seconds and they were subsequently centrifuged at 4,500 rpm for 3 minutes using a centrifuge (MX-305; Tomy, Co., Ltd., Tokyo) to precipitate the remains of spider bodies. 50 μl supernatant of each sample solution was then mixed with 0.6 μl proteinase K solution (Takara Bio Inc., Kusatsu) and 9.4 μl of the lysis buffer mentioned above. The samples were then processes under the temperature conditions of 37 °C for 60 min followed by 95 °C for 10 min.

### PCR and Illumina sequencing

For the DNA metabarcoding of prey Hexapoda, we used the Chiar16SF-Chiar16SR^49^ primer pair targeting the mitochondrial 16S rRNA region of Hexapoda. In general, detection of prey Hexapoda DNA from spider bodies is hampered by the non-selective PCR amplification of spider sequences. Even if Hexapoda-specific primers are used with blocking primers targeting spider DNA, the ratios of prey sequences to spider sequences in DNA metabarcoding data are very low^50^. After performing several rounds of preliminary DNA metabarcoding assays using previously reported protocols, we found that the use of the Chiar16SF-Chiar16SR primer pair^49^ could yield best results of prey detection from spider bodies.

For each sample, the amplification of the mitochondrial 16S rRNA region was performed with the forward primer Chiar16SF fused with 3-6-mer Ns for improved Illumina sequencing quality^51^ and the forward Illumina sequencing primer (5’ - TCG TCG GCA GCG TCA GAT GTG TAT AAG AGA CAG- [3–6-mer Ns] – [Chiar16SF] -3’) and the reverse primer Chiar16SR^49^ fused with 3–6-mer Ns and the reverse sequencing primer (5’-GTC TCG TGG GCT CGG AGA TGT GTA TAA GAG ACA G [3–6-mer Ns] - [Chiar16SR] -3’) (0.2 μM each). The PCR reaction was conducted using DNA polymerase system of KOD One (Toyobo Co. Ltd., Osaka). The temperature profile of the PCR was by 40 cycles of 98 °C (denaturation) for 10 s, 54 °C (annealing) for 5 s, and 68 °C (extension) for 30 s, and final extension at 68 °C for 2 min. The DNA extraction products were centrifuged at 10,000 rpm for 5 min before each PCR reaction not to carry PCR inhibitor chemicals potentially remained in the template DNA solutions. To prevent generation of chimeric sequences, the ramp rate through the thermal cycles was set to 1 °C/sec^52^. Illumina sequencing adaptors were then added to respective samples in the supplemental PCR using the forward fusion primers consisting of the P5 Illumina adaptor, 8-mer indexes for sample identification^53^ and a partial sequence of the sequencing primer (5’-AAT GAT ACG GCG ACC ACC GAG ATC TAC AC - [8-mer index] - TCG TCG GCA GCG TC -3’) (0.43 μM each) and the reverse fusion primers consisting of the P7 adaptor, 8-mer indexes, and a partial sequence of the sequencing primer (5’-CAA GCA GAA GAC GGC ATA CGA GAT - [8-mer index] - GTC TCG TGG GCT CGG -3’) (0.43 μM each). KOD One was used with a temperature profile of 98 °C for 10 s, followed by 8 cycles of 98 °C for 10 s, 55 °C for 5 s, and 68 °C for 30 s (ramp rate = 1°C/s), and final extension at 68°C for 2 min.

The PCR amplicons of the samples were then pooled after a purification/equalization process with the AMPureXP Kit (Beckman Coulter, Inc, Brea). Primer dimers, which were shorter than 200 bp, were removed from the pooled library by supplemental purification with AMpureXP: the ratio of AMPureXP reagent to the pooled library was set to 0.6 (v/v) in this process. The sequencing libraries of the mitochondrial 16S rRNA region was processed in an Illumina MiSeq sequencer (run center: KYOTO-HE; 15% PhiX spike-in). Because the quality of forward sequences is generally higher than that of reverse sequences in Illumina sequencing, we optimized the MiSeq run setting in order to use only forward sequences^54^. Specifically, the run length was set 271 forward (R1) and 31 reverse (R4) cycles to enhance forward sequencing data: the reverse sequences were used only for screening mitochondrial 16S rRNA sequences in the following bioinformatic pipeline.

### Bioinformatics

The raw sequencing data were converted into FASTQ files using the program bcl2fastq 1.8.4 distributed by Illumina. The output FASTQ files were demultiplexed with the program Claident v0.2. 2018.05.29^55^, by which sequencing reads whose 8-mer index positions included nucleotides with low (< 30) quality scores were removed. Only forward sequences were used in the following analyses after removing low quality 3’ -ends using Claident. Noisy reads^55^ were subsequently discarded, and subsequently, potentially chimeric reads were removed as well using the programs UCHIME^56,57^ (*de novo* mode) as implemented in Claident. The filtered reads were clustered with a cut off sequencing similarity 97% using the program VSEARCH (Rognes et al., 2014). The operational taxonomic units (OTUs) representing less than 10 sequencing reads were subsequently discarded. The molecular identification of the remaining OTUs was performed based on the combination of the query-centric auto-*k*-nearest neighbor (QCauto) method^58^ and the lowest common ancestor (LCA) algorithm^59^ as applied using Claident (reference DNA database version, v0.9.2021.3.25). On average, 3,140.7 spider reads (SD = 1,844.8) and 466.0 Hexapoda reads (SD = 846.5) were obtained from each sample. The spider sequences were used to supplement identification of spider species (in particular, identification of juvenile spiders), while Hexapoda sequences were used for the analysis of prey compositions. Although potential predation events between spiders might be detectable with the spider sequences, the use of the Hexapoda-targeting primers could result in unexpected biases. Therefore, we did not use the spider data for the inference of spider–spider interactions: developing DNA metabarcoding protocols for intra-guild predation of spiders require future intensive studies.

Based on the DNA sequence data, a “sample-level” matrix (Araneae × Hexapoda matrix) representing the number of sequencing reads of respective Hexapoda OTUs in respective spider samples, was obtained for each month. Based on the matrix of each month, a binary sample-level matrix, which indicated whether each OTU was present or absent in each sample, was further obtained: prey OTUs with 10 or more sequencing reads were designated as present (1) in a spider sample, while those with less than 10 reads were designated as absent (= 0) because they might result from unexpected contamination events during sampling, DNA extraction, or PCR. The binary sample-level matrices of April–November datasets, in total, included 50 spider species and 974 Hexapoda OTUs. At least one Hexapoda OTU was detected from 1,556 out of 2,224 spider samples: i.e., potential prey information was obtained from 70.0 % of the spider specimens examined. Based on the binary sample-level matrices, we further obtained “species-level” matrices, in which rows represented spider species, columns depicted Hexapoda OTUs, and cell entries indicated the number of samples from which respective spider–Hexapoda OTU combination were observed (hereafter, prey detection count).

### Predator and prey diversity

The species compositions of sampled spiders were visualized as a bar graph across the months. Likewise, the family-level taxonomic compositions of prey were shown as a bar graph based on the prey detection counts of the species-level matrices. The compositions of prey ecological guilds were visualized as well. In the analysis, prey OTUs were classified into four categories, namely, detritivores, herbivores, predators/parasites, and organisms with variable feeding habits, based on the reference search of family-level taxonomy. It should be acknowledged that each family possibly included species belonging to different guilds. Therefore, the list is an incomplete proxy based on currently available information, requiring careful interpretation in its use. The classification of guilds should be continually updated based on lower-level taxonomic information in future studies.

To examine how prey compositions varied across spider species and sampling months, we constructed a permutational multivariate analysis of variance^60^ (PERMANOVA) implemented in the R vegan package (9,999 permutations). The family-level composition of prey detection counts was used as the response variable, while spider species, sampling months, and interactions between them were set as explanatory variables.

### Network structure

Based on the species-level Araneae × Hexapoda matrices, networks depicting potential trophic interactions between spiders and Hexapoda were visualized for respective months with the R bipartite 2.1.6 package^61^. To evaluate seasonal changes in network structure, a series of network topological indices were calculated. For each month, we calculated the checkerboard scores representing the avoidance of prey/predator ranges within spider/prey communities, *H2’* metric of community-level interaction specialization, weighted NODF metric of nestedness, and Barber’s modularity index of network compartmentalization^34^. To quantify the extent to which the observed network pattern was deviated from that expected by chance, each of the network indices was standardized based on a randomization analysis as follows^33^:

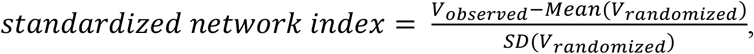

where *V*_*observed*_ represented the observed value of an applied network index, and *mean*(*V*_*randomized*_) and *SD*(*V*_*randomized*_) were the mean and standard deviation of the index values of randomized matrices, respectively. To make randomized matrices, the labels of spider species were shuffled within each sample-level matrix^33^ (1,000 permutations). The randomized sample-level matrices were converted into species-level matrices for the calculation of network indices. In the calculation of checkerboard scores, *H2’*, and weighted NODF nestedness, the bipartite package of R was used. Barber’s metric of network modularity was calculated using the program MODULAR^62^. Significance of the estimates was examined with Benjamini-Hochberg adjustment of *P*-values the based on the randomization [false discovery rate (FDR) < 0.05].

### Genealogy analysis of network modules

Based on the species-level matrix of each month, spider species and prey OTUs were grouped into network modules of closely interacting vertices (species/OTUs) using the infomap algorithm^63^. Then, the “genealogy” of network modules across months was estimated with the alluvial diagram (flow diagram) approach^64^ using the Alluvial Diagram Generator^65^. The diagram indicated the flow of module components (spider species and prey OTUs) succeeded across different time points.

### Interaction generality and network centrality

To quantify the extent to which each spider species or prey OTU interacted specifically to its prey or predators, we calculated the *d’* index of standardized Kullback-Leibler distance for each month using the R bipartite package. Because *d’* score varies from 0 (minimum specificity given a species abundance within the community) to 1 (maximum specificity given a species abundance), the value 1 – *d’* was used as a measure of the extent to which a spider species or prey OTU interacted with broad ranges of prey/predators (“interaction generality”).

To evaluate the extent to which each species/OTU was located at the core position within each network, we also calculated betweenness network centrality. Betweenness centrality is the measure of the degree to which a given vertex is located within the shortest paths connecting pairs of other vertices in a network. Scores of betweenness were normalized for each network so that they varied from 0 (occupation at marginal positions within a network) to 1 (occupation at shortest paths for all pairs of vertices) using the igraph v.1.3.0 packag^66^e of R.

These indices are expected to help us highlight species or OTUs with potentially great impacts on community-level processes. Considering the history of discussion on such species^9,10^, we called them “potential” keystones. In community ecology, a keystone species is defined as a species with disproportionately large ecological influence on surrounding biotic and abiotic environments relative to its abundance^9,10^. The interaction generality index is expected to highlight predator species having disproportionately large impacts on diverse prey populations relative to their abundance as well as prey OTUs having disproportionately large influence on predator populations relative to their abundance. Meanwhile, in terms of impacts on whole-community processes, impacts of a species may be negligible if its population size or biomass is extremely small. Such rare species/OTUs are given low scores in the metric of betweenness centrality. Thus, we evaluated species on two dimensional surfaces consisting of interaction generality and betweenness centrality. Removal of species simultaneously showing high interaction generality and high betweenness centrality are expected to cause breakdown of predator–prey interaction networks, imposing great impacts on ecosystem-level structure and functions.

### Meta-network analysis

To infer the structure of the meta-network representing all the interactions observed from April to November, we compiled the eight species-level matrices representing predator–prey interactions observed in respective months. Modules within the meta-network were detected based on the infomap algorithm mentioned above. The number of links to other vertices within the same module was counted for each spider species or prey OTU: the within-module degree was *z*-standardized (zero-mean, unit-variance). We also calculated among-module connectivity, which represented how a vertex is positioned in its own module and with respect to other modules^37^. By definition, among-module connectivity could vary between 0 (all links are within the module a target vertex belongs to) and 1 (links are distributed uniformly among all modules). Albeit somewhat arbitrary, species with *z*-standardized within-module degree greater than 2.5 and among-module connectivity greater than 0.62 have been designated as “super generalists” in studies of ecological networks^36^.

## Data availability

The 16S rRNA sequencing data are available from the DNA Data Bank of Japan (DDBJ) with the accession number PRJDB12701. The microbial community data are deposited at our GitHub repository (https://github.com/hiro-toju/Spider_prey_networks.git) [to be publicly released after acceptance of the paper].

## Code availability

All the scripts used to analyze the data are available at the GitHub repository (https://github.com/hiro-toju/Spider_prey_networks.git) [to be publicly released after acceptance of the paper].

## Acknowledgements

This work was financially supported by JSPS Grant-in-Aid for Scientific Research (18H04009) and JST FOREST (JPMJFR2048) to H.T..

## Author Contributions

H.T. designed the work with S.S.. S.S. performed experiments. S.S. analyzed the data with H.T.. S.S. and H.T. wrote the paper with Y.G.B..

## Additional information

**Supplementary information** is available for this paper at [URL to be supplied by the publisher]

## Extended Data Figure legends

**Extended Data Fig. 1** | **Order- and genus-level taxonomy of detected prey. a**, Order-level analysis. The compositions of prey detection counts are shown across the sampling months in terms of the order-level taxonomy. **b**, Genus-level analysis. The compositions of prey detection counts are shown across the sampling months in terms of the genus-level taxonomy. **c**, Effects of spider species and sampling months on prey compositions. A PERMANOVA of family-level prey compositions (Fig. 1b) was performed by setting spider species, sampling months, and interactions between them as explanatory variables.

**Extended Data Fig. 2** | **Counts of prey detection from respective spider species**. The compositions of prey detection counts are shown for each spider species across the sampling months. Top 19 spider species with highest numbers of specimens with prey sequences are presented.

**Extended Data Fig. 3** | **Hunting type of spiders**. For each category (hunting type) of spiders (as defined in Fig. 2), the compositions of prey detection counts are shown with putative ecological guilds of prey. Based on family-level taxonomy, the prey guilds were classified into four categories: detritivores, predators/parasites, herbivores, and omnivores (variable feeding habits). The prey OTUs unidentified at the family level were omitted in the graphs.

**Extended Data Fig. 4** | **Network topology**. Within the networks depicting the predator–prey interactions observed in respective months, the family-level taxonomy of prey vertices is indicated. The layout of the vertices was optimized based on the “stress” algorithm of network ordination as implemented in the ggraph package of R.

**Extended Data Fig. 5** | **Network modules**. Within predator–prey interaction network of each month, vertices were classified into modules consisting of closely interacting spiders and prey based on the infomap algorithm.

**Extended Data Fig. 5** | **Temporal variability in topological roles**. For the three spider species collected throughout the seasons (spider species that appeared in seven or eight months), temporal shifts of interaction generality and betweenness network centrality are presented. The size of symbols is roughly proportional to the number of spider individuals analyzed. Color indicates sampling months.

