## Extended Data Fig. 1 for "Dynamics of species-rich predator–prey networks and seasonal alternations of keystone species"

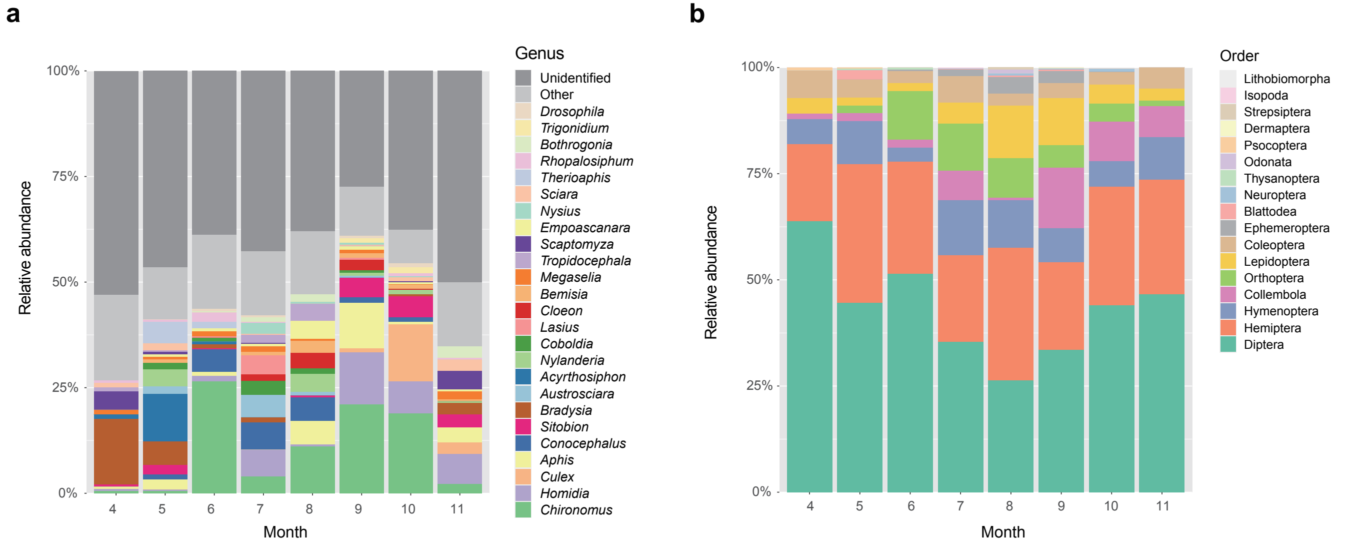


**c**

| Explanatory variable | df | $F_{model}$ | $R^2$ | $P$ |
|---|---|---|---|---|
| Spider species | 49 | 2.99 | 0.077 | 0.0001 |
| Month | 7 | 27.93 | 0.103 | 0.0001 |
| Spider species × month | 118 | 1.53 | 0.095 | 0.0001 |

Response variable = prey compositions (family level)
