## Supplementary figures and images for "Dynamics of species-rich predator–prey networks and seasonal alternations of keystone species"

### Extended Data Fig. 2

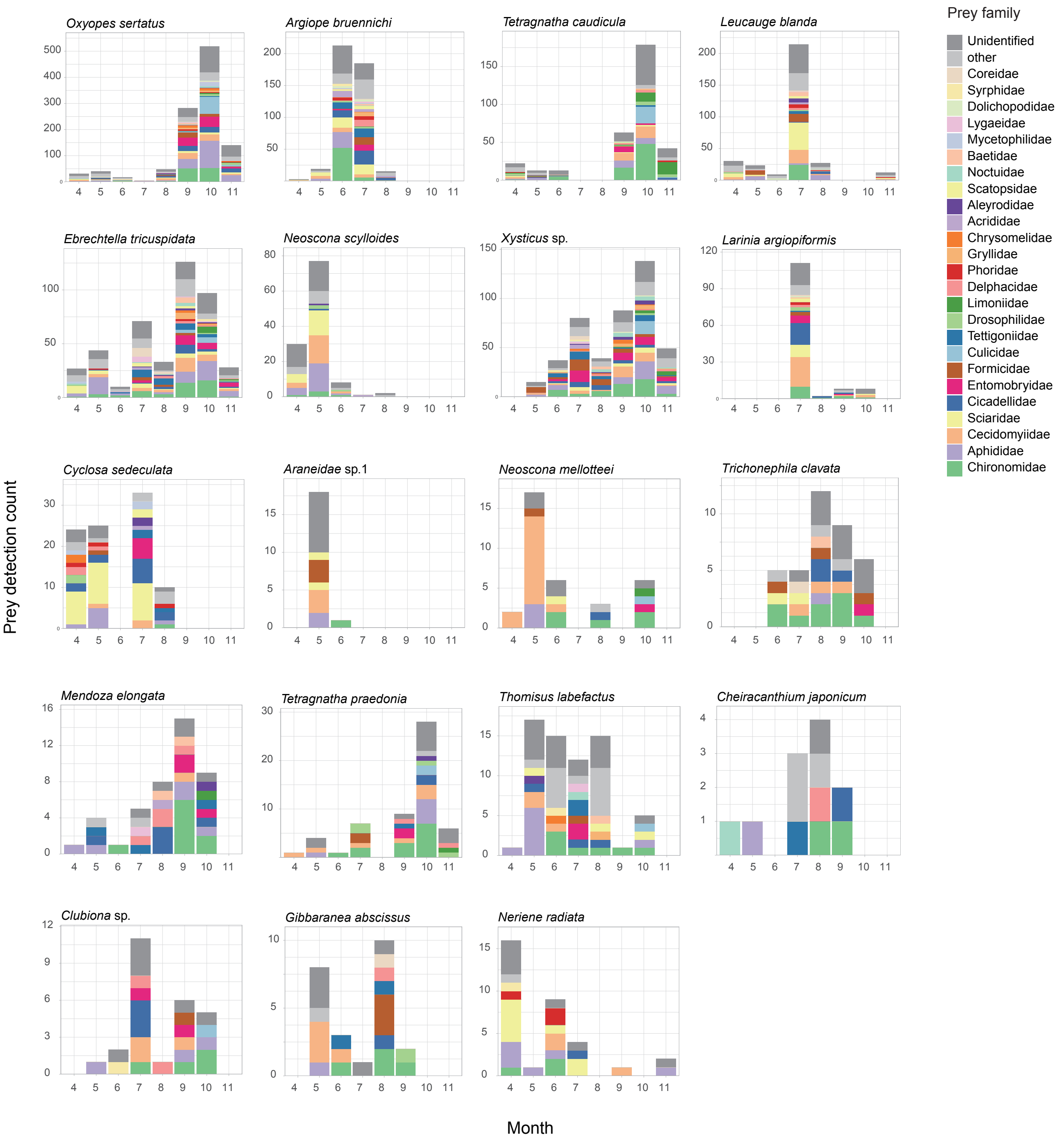

### Extended Data Fig. 3

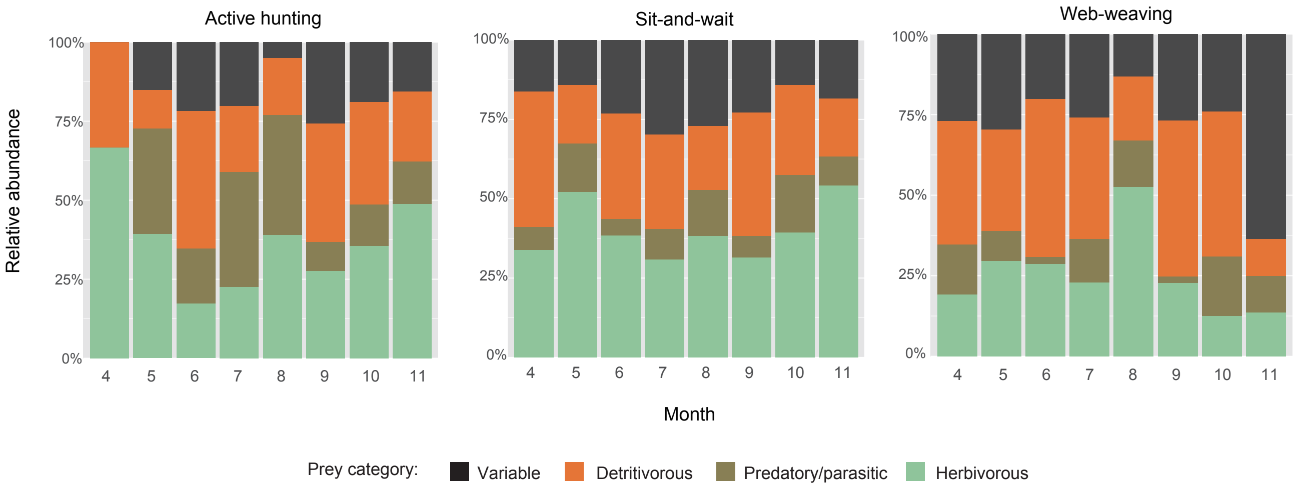

### Extended Data Fig. 4

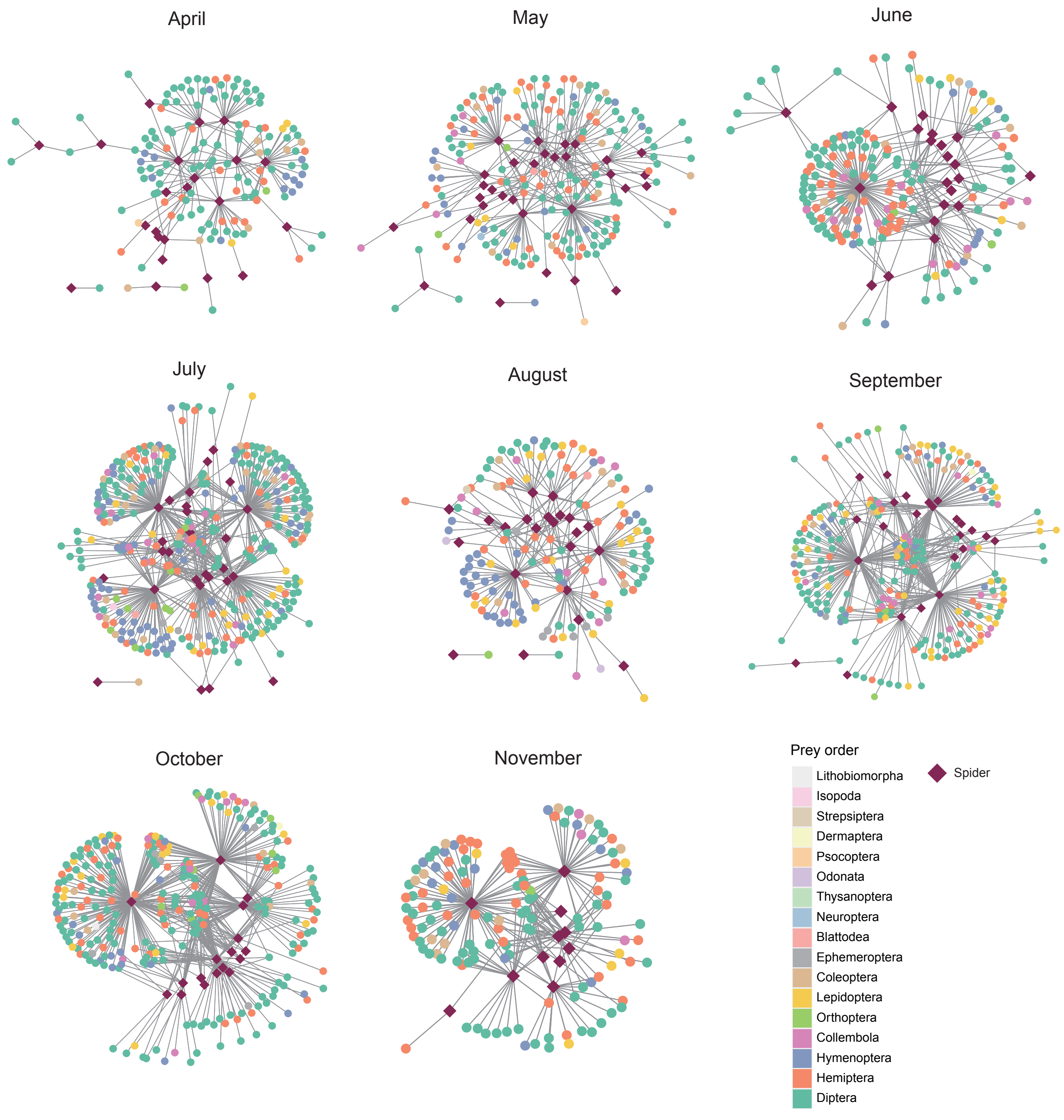

### Extended Data Fig. 5

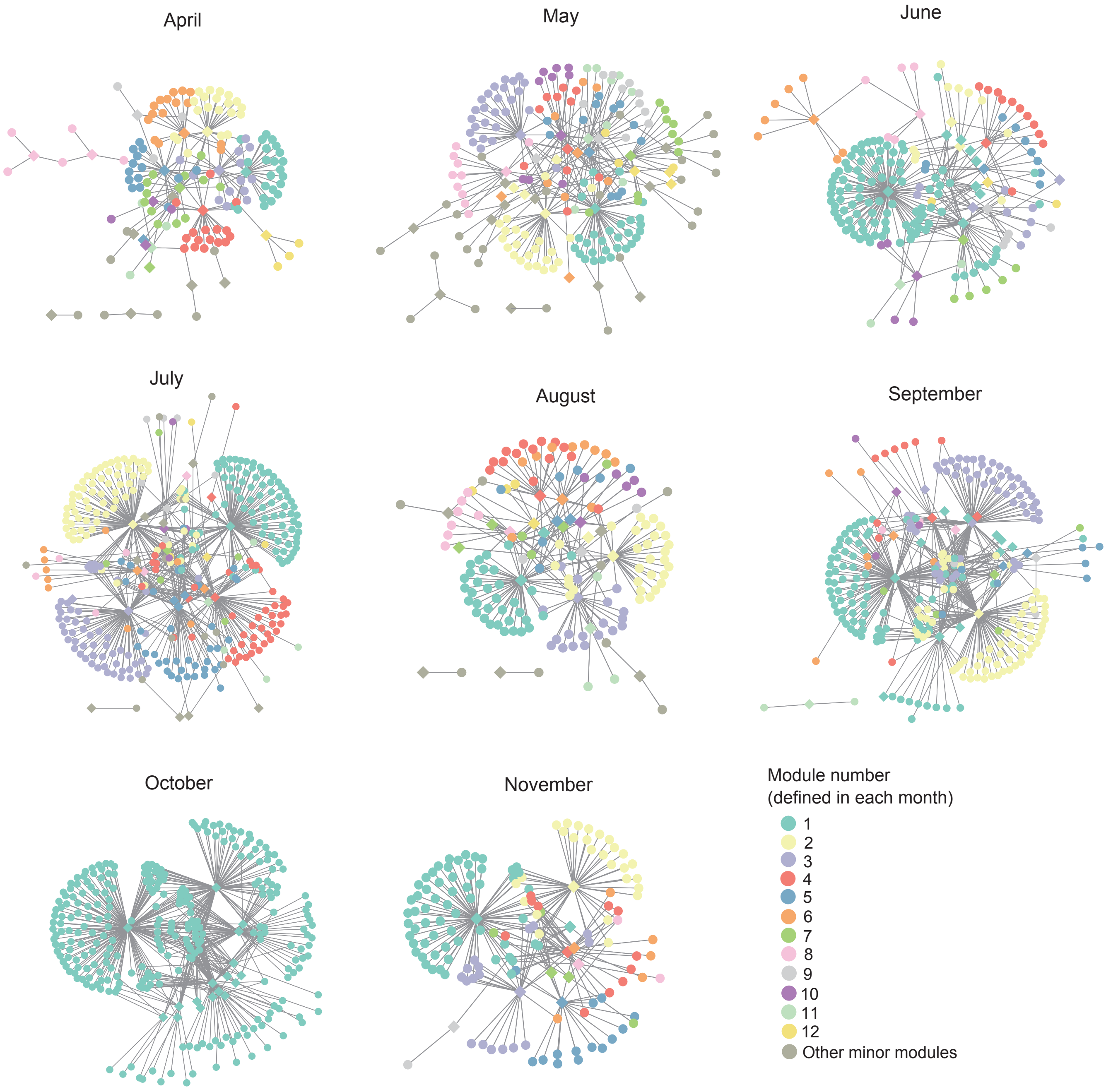

### Extended Data Fig. 6

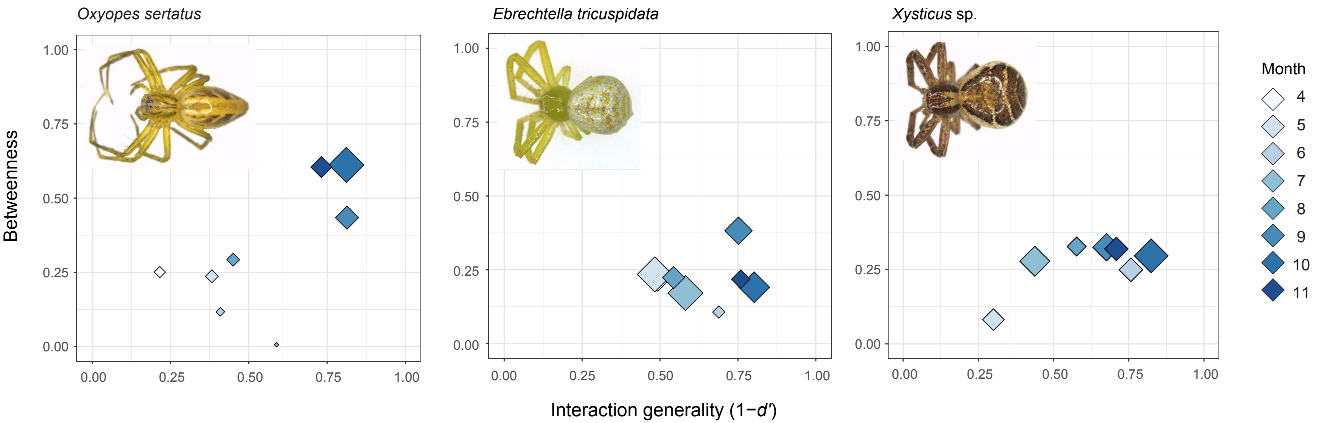
